# Formation of membrane invaginations by curvature-inducing peripheral proteins: free energy profiles, kinetics, and membrane-mediated effects

**DOI:** 10.1101/2022.11.09.515891

**Authors:** Mohsen Sadeghi

## Abstract

Peripheral proteins, known to induce curvature, have been identified as key agents in the spontaneous remodeling of bilayer membranes, leading to invaginations and the genesis of membrane tubules. For proteins like cholera and Shiga toxin, which impart the membrane with locally isotropic curvatures, the resultant membrane-mediated interactions remain notably subtle. Consequently, the collective action of these proteins, culminating in the formation of dense clusters on the membrane and subsequent invagination, unfolds over an extended timescale, often spanning several minutes. This gradual progression challenges direct simulation of the invagination process, even with coarsegrained models. In this study, we introduce a steered molecular dynamics protocol wherein peripheral proteins are impelled to converge on a membrane patch, instigating an invagination. Utilizing the Jarzynski equality, we derive the free energy profile of this process from a suite of non-equilibrium simulation replicas. Examining two distinct membrane-associated proteins, we elucidate the influence of protein flexibility and the distribution of induced curvatures on both the remodeling process and the corresponding free energy profile. We delve into the role of membrane-mediated effects in shaping protein organization within the invaginated domain. Building on the free energy profile, we model the formation of invaginations as a Markovian process, and offer estimates of the corresponding timescales. Our findings yield minute-long implied timescales that resonate well with empirical observations.

## I. INTRODUCTION

Living cells harness a myriad of mechanisms to transport materials to and from their surroundings [1]. While small molecules and ions can either diffuse directly across the membrane [2] or utilize channel proteins [3], most larger entities–including biomolecular assemblies and virions–primarily depend on endo/exocytosis trafficking (few exceptions with unconventional pathways notwithstanding [4]). This process entails significant membrane remodeling, in the form of vesicular or tubular intermediates. Key mediators of endocytosis include scaffolding proteins like clathrin [5], caveolin [6], and endophilin [7]. Additionally, active cellular machinery, exemplified by actin and dynamin assemblies, often plays a pivotal role by directly applying force to the membrane [8, 9]. Finally, liquid-like protein condensates on the membrane have been suggested to contribute to endocytosis [10, 11]. Conversely, certain toxins, such as Shiga, cholera, and ricin, internalize via tubular invaginations, bypassing both mechanisms [12–15]. Both cholera and Shiga toxin possess distinct pathogenic and membrane-bending subunits. The latter, when densely clustered, can cooperatively bend the membrane into extended tubules, facilitating the internalization of the pathogenic entities—a process encapsulated by the glycolipid-lectin (GL-Lect) hypothesis [16, 17]. Beyond its toxicological implications, this mechanism presents captivating prospects for drug or antigen delivery into living cells [17].

Tubular membrane invaginations can spontaneously arise through a variety of physical mechanisms. A prominent example is the adsorption of molecules that alter the membrane’s spontaneous curvature, subsequently inducing a tension that favors tubulation [18]. Protein-protein crowding is another significant factor driving membrane tubulation [19]. Membrane compression, whether due to osmotic pressure-induced volume reduction or an increase in membrane area, can foster the development of stable membrane tubules [20]. This is analogous to a mechanism recently proposed for the swift formation of endocytic pits in the synaptic cleft [21]. For further examples, we recommend ref. [22] and the references therein.

To delve into the progression of membrane budding and tubulation influenced by membrane-bending peripheral proteins, evaluating the free energy landscape of this process is paramount. As was recently demonstrated by Groza et al., through experiments with controlled binding affinities of multivalent globular binders, the binding energy is the determining factor in the success of these membrane-bending constructs to be spontaneously endocytosed [23]. Ewers et al. proposed a theoretical model that juxtaposed the local curvature induced (and favored) by each protein and its associated energy against the substantial energy barrier posed by the initiation of tubular invagination. This model also considered the balance between energy gained from species joining the tubule and the entropy lost from their concentration, analyzing the free energy profile for spontaneous tubule nucleation in various scenarios concerning toxins and virions [24]. Through Monte Carlo simulations, Šarić and Cacciuto quantified the free energy landscape associated with tube formation when multiple large colloidal particles bind to the membrane, based on varying particle-particle separations. Their findings underscored the preference for tubular invagination over other budding configurations [25]. In another approach, Tourdot et al. employed a continuum membrane model alongside curvature fields, tracking changes in the excess chemical potential of membrane-bending proteins to pinpoint the tubulation threshold [26]. Mathijs et al. adopted a Ginzburg-Landau-style free energy for the protein coat, examining the stability of cylindrical tubes adorned with proteins [27]. Furthermore, Mahapatra and Rangamani constructed a theoretical framework estimating the free energy of membrane tubulation resulting from bound BAR-domain proteins exhibiting anisotropic curvature. Their model highlighted the distinct snap-through properties observed during dome-to-cylinder transitions [28]. Although these studies shed considerable light on the topic, they often sidestep the dynamic interplay between membrane and proteins, subsequently offering a constrained perspective on the free energy and kinetics of tubulation.

In this work, we present a novel perspective on membrane invagination using a mesoscopic dynamical membrane model [29]. Analyzing membrane invagination influenced by a collection of peripheral proteins poses inherent scaling challenges. While all-atom or bottom-up coarse-grained models like MARTINI [30] offer detailed insights into protein-membrane interactions, composition variability, protein flexibility, and membrane dynamics, they suffer from restrictive spatiotemporal scopes. Conversely, continuum models provide expansive scale ranges, but sacrifice many such intricate details. Attempts to amalgamate the strengths of both models have been made using multiscale approaches [31, 32]. Despite their utility, these strategies often restrict the transfer of information contained in each scale to heuristically selected features, risking the omission of local effects that might play a global role.

Our approach adopts a single-scale strategy, utilizing a mesoscopic model infused with richer information than typical membrane models of its kind. This model has been demonstrated to mimic realistic kinetics associated with both equilibrium and non-equilibrium membrane fluctuations [33, 34] as well as the lateral diffusion of membrane-bound peripheral proteins [35]. Importantly, it naturally replicates the complex membrane-mediated interactions resulting in protein aggregation [35, 36], and facilitates intricate calculations of free energy and entropy, shedding light on the forces at play [35, 37]. Finally, it comfortably fits in meso-scale models of large biological systems [33, 38]. Positioning itself midway between scale and detail, our model captures kinetics of the proposed pathway and fluctuation effects, but also faces time constraints in unbiased simulations.

To address this limitation while leveraging the model’s dynamical nature, we introduce a scheme for steered simulations. Here, peripheral proteins are deliberately guided along the invagination pathway. Invoking the renowned Jarzynski equality, we extract the free energy profile throughout the process, sidestepping the need for equilibrium sampling. We further assess the kinetics of its various phases using transfer operator theory. Engaging two distinct protein candidates, differing in their membrane curvature inductions, we illuminate how a switch between them intricately influences the free energy landscape, resultant membrane structures, and kinetics.

## RESULTS

### Modeling membrane-bending peripheral proteins

To have a computationally efficient and yet dynamically reliable model of membrane and bound peripheral proteins, we have employed the framework previously introduced by the author and collaborators [29, 33, 35, 42] (see Methods section “Simulations” for more details). Peripheral proteins, capable of inducing local membrane curvature, are incorporated into the model using force field masking (Fig. 1A). This entails using modified as well as reinforcing local contributions to the potential function. The protein particles are nevertheless allowed to diffuse laterally, with accompanying force field-modifying moves handled in a Monte Carlo scheme [35, 42]. An elastic protein model, characterized by intrinsic curvature and finite flexibility, in interplay with the underlying membrane, is employed to parameterize the masked region. In our simulations, two distinct protein models are used, both featuring an intrinsic curvature of 0.08 nm*−*1, but with different flexibilities (Methods section “Simulations”). Throughout this work, these two protein models are referred to as MP_1_ and MP_2_.

**Figure 1:**
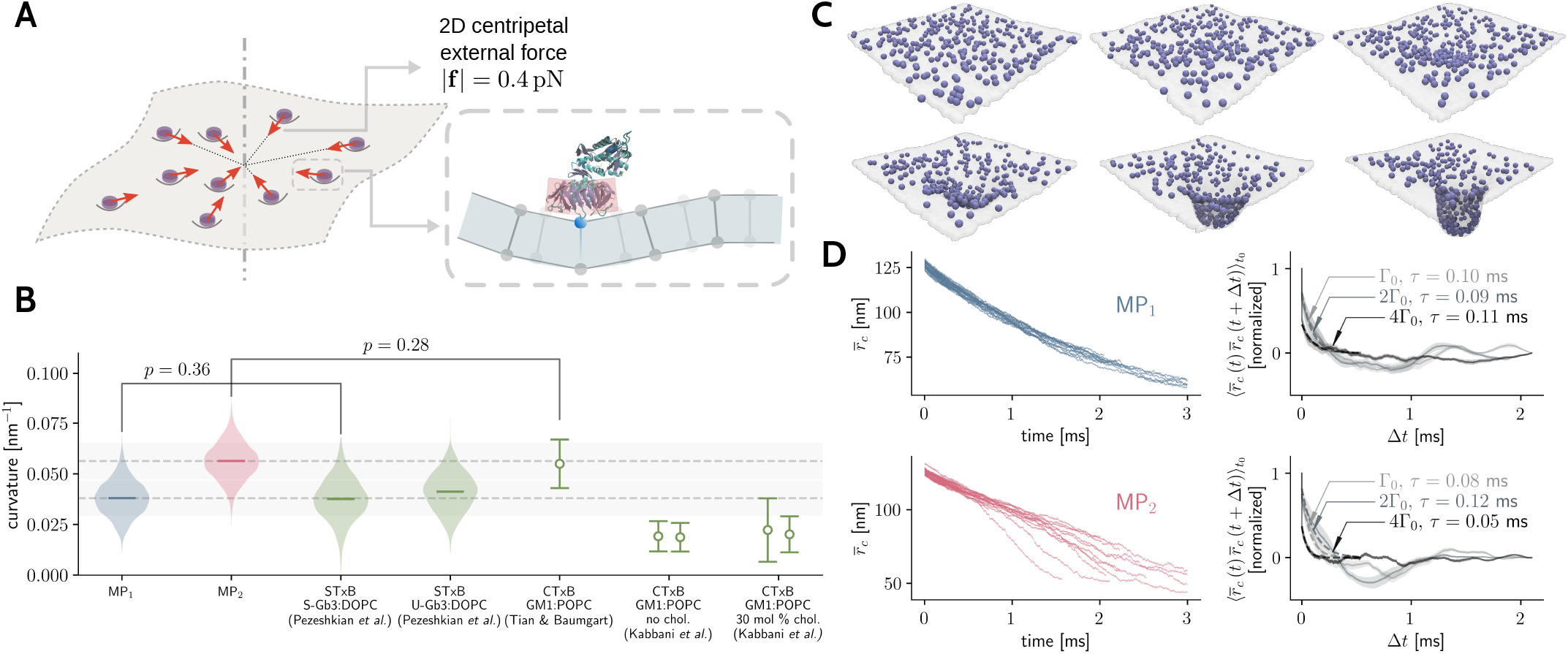
Characterizing the formation of membrane invaginations through steered simulations with membrane-bending peripheral proteins: **(A)** A schematic illustration of the simulation setup: peripheral proteins are dispersed on the membrane and drawn toward the patch’s center via in-plane centripetal external forces. An inset provides a closer look at a curvature-inducing peripheral protein on the particle-based membrane model. **(B)** Distribution of the local membrane curvatures generated by the protein types, MP_1_ and MP_2_. The curvature data are compared with findings from Pezeshkian et al. [39], Tian and Baumgart [40], and Kabbani et al. [41]. Gray bands and error bars correspond to standard deviations. p-values are the result of Mann-Whitney U tests, with larger values implying more similar distributions. **(C)** A sequence of snapshots from a single instance of the steered simulations (see Supplementary Movies). **(D)** (left) presentation of mean in-plane radial distances of proteins, measured from the central axis of the membrane patch and denoted as 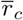, over time during the steered simulations. Data from different simulation replicas are included. (right) the time-correlation of the mean in-plane radial distance from a set of unbiased simulations (sans external forces) with the same proteins as denoted on the left. Unbiased simulations are conducted at the same surface concentration as the steered counterparts (Γ_0_), as well as higher concentrations of 2Γ_0_ and 4Γ_0_. Timescales derived from an exponential decay model accompany each dataset.

Initially, we conducted unbiased simulations with both protein models on tensionless membranes to determine, among other properties, the distribution of induced local curvatures. Fig. 1B showcases these distributions and juxtaposes them against: (i) Tian and Baumgart’s curvature measurements for cholera toxin subunit B (CTxB) on POPC membranes, via tether pulling and observing protein sorting [40], (ii) curvature assessments by Kabbani et al. related to CTxB’s influence on POPC membranes with and without cholesterol, by measuring the radii of buds induced by CTxB’s [41], and (iii) molecular dynamics simulations by Pezeshkian et al., detailing curvature estimates for Shiga toxin subunit B (STxB) on DOPC membranes containing Gb3 receptors, with saturated or unsaturated acyl chains [39]. We numerically contrasted our estimated curvature distributions with (i) and (iii) using the Mann-Whitney U non-parametric test [43]. Resulting p-values highlight the resemblance of MP_1_ to STxB from Pezeshkian’s simulations and MP_2_ to CTxB measurements by Tian and Baumgart (Fig. 1B). This comparison helps ground the two protein models, although the variance in the reference data could be inherent to the proteins and membrane compositions or characteristic to the measurement techniques.

### Steered simulation of the invagination pathway

At the outset of each simulation, MP_1_ or MP_2_ peripheral proteins are randomly distributed across the membrane’s surface. An external centripetal force with a magnitude of 0.4 pN is exerted on each protein particle (Fig. 1A). Influenced by these forces, the proteins eventually aggregate at the membrane patch’s center, forming a pit, which may subsequently develop into an invagination (Fig. 1C and Supplementary Movies).

In this study, we adopt the mean in-plane radial distance of protein particles relative to the simulation box’s central axis as the reaction coordinate describing the pit formation and invagination (eq. (5)). The time evolution of this reaction coordinate, denoted 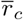, for all simulation replicas are shown in Fig. 1D. These simulations amount to the total time of 38.4 ms with MP_1_ (31 replicas) and 32.1 ms with MP_2_ (30 replicas).

Before delving into the free energy estimation, it’s prudent to discern how slowly the steered simulations are executed and ascertain how far the membrane system deviates from equilibrium. One approach is to compare simulation durations to the system’s physical timescales. Out-of-plane membrane fluctuations, characterized by the wave vector **q**, have a hydrodynamic dissipation timescale computed as 4*η/κq*^3^, where *η* represents the solvent’s viscosity, and *κ ≈* 20 *kT* is the membrane’s bending rigidity [44]. This results in a hydrodynamic dissipation timescale of about 0.01 ms. One extra remark to this computation is that the assembly of membrane-bending proteins can diminish the membrane’s effective bending rigidity [45], consequently prolonging the hydrodynamic relaxation timescale. Nevertheless, in our context, this still ranks significantly below other system timescales explored subsequently.

With the current model, and via unbiased simulations, we have determined both protein to have a lateral diffusion coefficient of *D*_p_ = 0.30 *±* 0.03 μm^2^ s^*−*1^ (following the same procedure laid out in [35]). This value compares well with the 0.35 *±* 0.09 μm^2^ s^*−*1^ estimate via fluorescence recovery after photobleaching (FRAP) assays on CTxB [40], or the 0.33 *±* 0.40 μm^2^ s^*−*1^ value, calculated from single-particle tracking trajectories of CTxB [41].

The typical duration required for proteins to traverse the average distance Δ*r* between adjacent neighbors can be estimated as Δ*r*^2^/4*D*_p_. Given the protein surface densities in our simulations, we surmise an upper bound of 0.1 ms for protein diffusion timescale. To further refine this estimate, we evaluated our unbiased simulations with a similar collective variable, 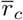, this time gauged against the center of mass of all the proteins distributed on the membrane patch. Examining the early decay in the time-correlation function of 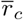 (Fig. 1D), we estimated timescales spanning between 0.05 ms and 0.11 ms, contingent on the protein variant and their surface concentration (Fig. 1D). Hence, in comparison, our steered simulations, spanning multiple milliseconds, transpire at a pace at least one order of magnitude slower than the system’s slowest process.

From another perspective, we can compare the magnitude of the external forces acting on proteins in our simulations with analogues from biological contexts. For instance, optical traps have been used to measure the unidirectional force exerted by the molecular motor kinesin to transport cargo along microtubules, revealing it can peak at 6.5 pN [46]. Another comparable force is that produced by a single actin filament’s growth, assessed at approximately 0.8 pN [47]. Consequently, the forces employed in our steered simulations are about an order of magnitude inferior to those exerted by motor proteins and roughly half of those imposed by a lone filament of the dynamic cytoskeleton.

Thus, while our free energy estimation protocol does not rely on equilibrium sampling, we have established that our steered simulations do no push the system too far off a quasi-equilibrium transition. Qualitatively, this means a rather wide variety of configurations are sampled with a close-to-canonical distribution in each step of the steered transition.

### Free energy profile of the invagination process

We move forward with utilizing the Jarzynski equation to discern the free energy profile associated with the formation of membrane invaginations [48, 49]. This method relies on sampling the external work done on the system in a collection of non-equilibrium transitions between macrostates (Methods section “free energy estimation” and eq. (1)). Fig. 2A illustrates the cumulative time series of external work measurements for the two proteins. While the work values span a broad range, the Jarzynski equation’s exponential weighting means that the majority of the weight is allocated to the smallest values. The free energy profiles are obtained by invoking eq. (1) between the scattered state with maximum 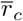 and the rest of states visited by the steered simulations (Fig. 2B).

**Figure 2:**
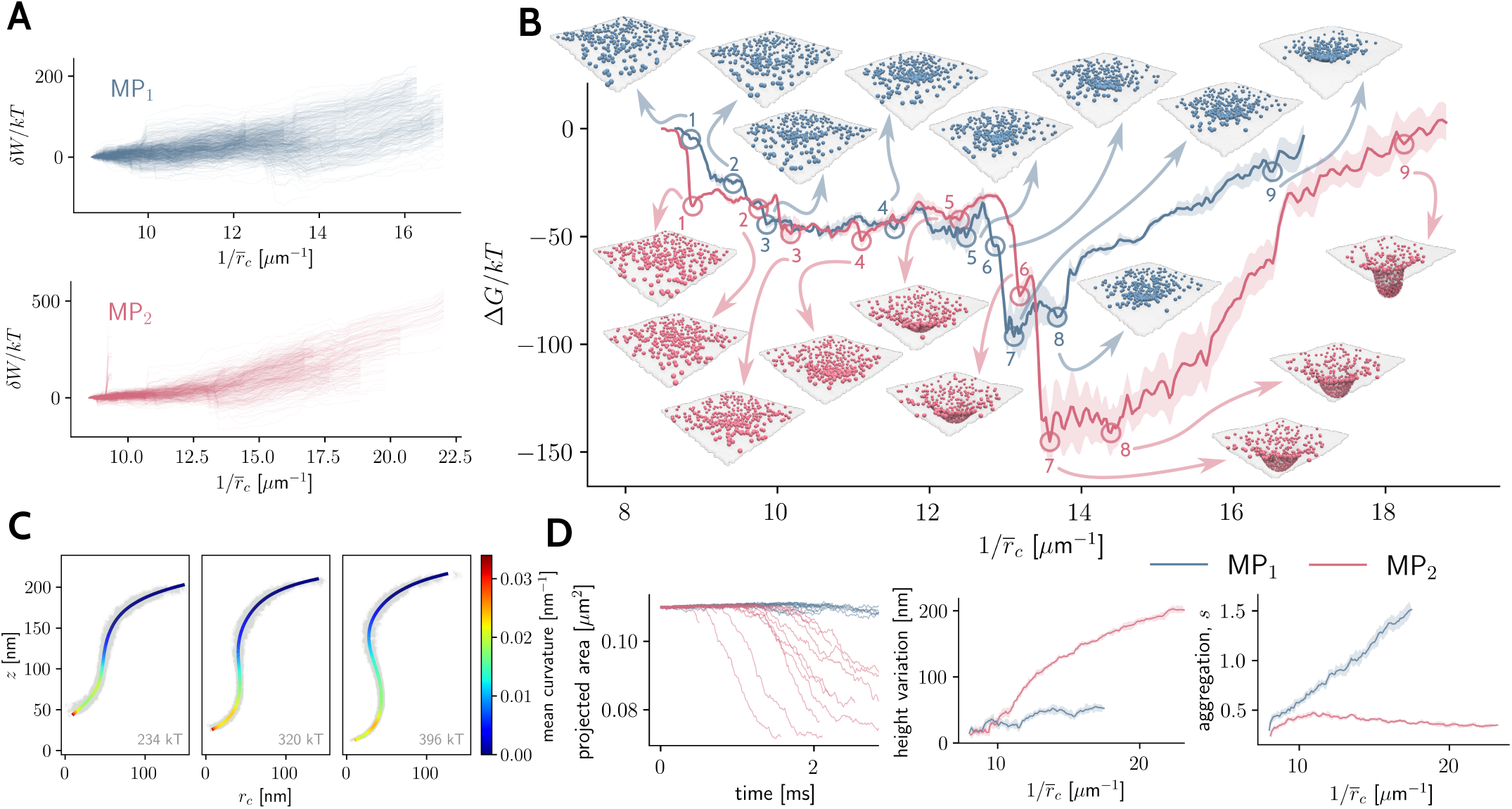
Exploring the free energy landscape of membrane invagination induced by peripheral proteins: Illustration of path-dependent external works for the proteins MP_1_ and MP_2_ derived from steered simulations. The work values are plotted against the inverse of the reaction coordinate, 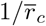. Thus, moving to the right on the graph amounts to visiting states with smaller 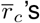, representing a sequence of aggregation, emergence of a central pit, and possibly invagination. **(B)** Depiction of the free energy landscapes for the two proteins as a function of the inverse reaction coordinate, illustrating aggregation and deformation. Free energy calculations are based on Jarzynski’s equation, with uncertainties estimated via bootstrapping; standard errors are displayed. Specific states on each curve are labeled for in-depth kinetic analysis. Accompanying each state, representative snapshots from a single steered simulation replica are provided. **(C)** Heatmap showcasing the positions of protein particles in the concluding phases of membrane invagination by MP_2_ proteins. This is recorded in a cylindrical coordinate system anchored on the optimal axis of symmetry (refer to the inset schematic in the far right). A polynomial curve of degree 7 is overlaid, with the colormap on the curve reflecting the mean curvature of the resulting surface of revolution. Each plot also presents an estimate of the membrane energy for the displayed region, obtained via the Helfrich model. **(D)** Various analytical plots pertaining to the investigated process, including: (left) time progression of the projected area of the membrane patch across different instances of steered simulations with both proteins. (middle) magnitude of fluctuations in the height of the membrane patch when viewed as a Monge patch, plotted against the inverse reaction coordinate. (right) the aggregation order parameter, denoted by *s*, presented for both protein types as a function of the inverse reaction coordinate.

We find the fact that the free energy profiles for both proteins have significant similarities quite noteworthy (Fig. 2B, consider the region between states 2 and 5). This serves both as a sanity check for the free energy estimation procedure, while simultaneously showcasing the fine differences due to the local curvature distributions associated with each protein.

In a broad perspective, the free energy profiles indicate that both proteins find the formation of a membrane pit or invagination favorable (Fig. 2B). For context, we can juxtapose these energy values against estimates relevant to the invagination process. This comparison can be facilitated by the Helfrich Hamiltonian: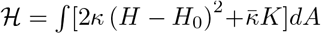, where parameters *H, H*_0_, and *K* represent the mean, spontaneous, and Gaussian curvatures, respectively, while *κ* and 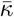 stand for the pertinent elastic constants [50, 51]. By employing the differential geometry of surfaces of revolution [52], we estimated the mean membrane curvature, as well as the membrane energy estimated by the Helfrich model, for an invagination induced by MP_2_ proteins during the final simulation phases, within the simulation replica visiting the smallest 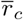(Fig. 2C).

Our analysis corroborates that the neck region typifies a minimal surface, in this context a catenoid, with a null mean curvature (Fig. 2C). Yet, regions populated by proteins typically exhibit a non-zero mean curvature. Leveraging the Helfrich functional, we discerned the total bending energy within the invaginated zone to be approximately 230 − 400 *kT* (Fig. 2C). As a point of comparison, morphing a flat membrane into a hemispherical cap demands an energy of 4*πκ*. Given *κ* = 20 *kT* [53], this translates to an energy of 251 *kT*. Meanwhile, a perfect cylindrical membrane tubule of length 150 nm and a mean tube radius of 50 nm – akin to our simulations – necessitates energy as elevated as 750 *kT*. The substantial energy prerequisite for membrane bending is mitigated by (i) the membrane acquiring a shape as close to a minimal surface as possible, and (ii) the coat of flexible proteins that alleviate the total elastic energy, especially when proximate to regions with curvatures close to their intrinsic curvature. Consequently, an energy dip materializes as proteins adeptly envelop the pits, counteracting the immense energies required for membrane deformation.

An evidence for this claim is to note the final gain in free energy between states 7 and 9 or beyond with MP_2_, which is *≈* 150 *kT* (Fig. 2B) and compare it with the similar increase in membrane energy (Fig. 2C). With the finite amount of proteins to coat the invagination in our simulations, the growth of the tubule is hindered, and further deformation leads to a comparable increase in the membrane energy and thus the free energy of the system.

Focusing on the early phases of the evolution along the reaction coordinate, there are marked dips in free energy for both proteins during the 1***→*** 2 and 2***→*** 3 transitions (Fig. 2B). These changes correspond to the creation of protein clusters on the membrane surface, with these clusters apparently more stable for MP_2_ (observed in states 1 and 3 in Fig. 2B). This evidence corroborates our earlier discoveries regarding cluster formation, where the more rigid proteins excelled in generating large clusters [35]. However, there may also exist another mechanism where extensive clusters of proteins, capable of strongly bending the membrane, globally repel one another due to membrane-mediated interactions of enthalpic origin [54, 55]. Such a phenomenon could account for the observed differences in the 2***→*** 3 or even 3***→*** 4 transitions between MP_1_ and MP_2_ proteins. With MP_1_, these transitions, which necessitate the congregation of smaller clusters at the center, are smoother and encompasse relatively minor barriers. Conversely, with MP_2_, they require surmounting substantial unevenness in the free energy landscape.

As previously mentioned, we identified significant free energy barriers at the onset of pit formation, notably in the 4 ***→*** 5 ***→*** 6 transitions (Fig. 2B). The energy required to overcome these barriers can be compared with the range of excess chemical potentials documented by Tourdot et al. in initiating membrane tubulation [26]. The conformation of the curved pit with the preferred curvature of the peripheral proteins, which reduces the total elastic energy, contrasts with the energetically unfavorable presence of proteins in the neck region, contributing to this barrier [14]. The spatial organization of dense protein clusters on the membrane may also be a contributing factor.

To delineate the disparities between MP_1_ and MP_2_ regarding their membrane deformation behaviors, we examined the membrane’s projected area (Fig. 2D). Given that the simulations were conducted for tension-less membranes coupled laterally to barostats (Methods section “Simulations”), area fluctuations are primarily attributed to the membrane’s out-of-plane deformations. Notably, while MP_1_ proteins induced a shallow pit in the membrane without a significant decrease in the projected area, MP_2_ proteins more dramatically reshaped the membrane, forming a deep invagination. Insight into these variations can also be gleaned by comparing the membrane height variations with the two proteins (Fig. 2D).

We discovered a substantial difference in how the particles of the two proteins are organized within the invaginated region. To quantify this, we employed an order parameter, defined as 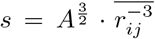, with *A* being the membrane projected area, *r*_*ij*_ the pairwise distance between protein particles, and the average taken over all pairs in one simulation frame. This parameter pertains to aggregation, as the rapid decay of 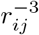 with pairwise distance ensures that the mean value correlates with the number of closely situated particles.

By plotting the order parameter *s* against the reaction coordinate 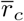, we unveiled that: (i) MP_2_ proteins generally preserve larger pairwise distances in all configurations, and (ii) while both MP_1_ and MP_2_ initially formed aggregates when forced toward the center, the latter quickly adopted a more dispersed configuration (Fig. 2D). The formation of invagination thus aids in achieving a more scattered distribution of MP_2_ proteins.

In sum, given the differences in free energy between the invaginated and flat states, we can assert that the formation of tubular invaginations, while a slow and energy barrier-driven process in the present context, is fundamentally irreversible. Reversion to previous states would necessitate alterations in other parameters, such as the membrane tension.

### Kinetics of the invagination process

To probe the kinetics underlying various stages of the invagination process, as illustrated with the free energy landscape, we employed a method rooted in transfer operator theory. This approach models the kinetics of an unbiased Markovian process in the 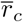 space (Methods section “Kinetics”). In essence, this approach relies on leveraging eq. (9) for finding transition probabilities in the presence of a potential of the mean force, depending on an estimate of the diffusion coefficient 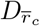.

Finding 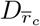, which is a collective parameter, is tantamount to understanding the friction encountered during transformation along the reaction coordinate. We anticipate that this friction is not constant but varies with 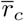. Our method involved a sliding window sub-sampling of trajectories along 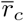, complemented by local calculations of mean squared displacements to ascertain diffusion coefficients. The outcome is a profile of average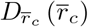, accompanied by uncertainties originating from finite sample sizes in each bin as well as inherent variations in microstates included in a partition along 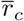(Fig. 3A).

**Figure 3:**
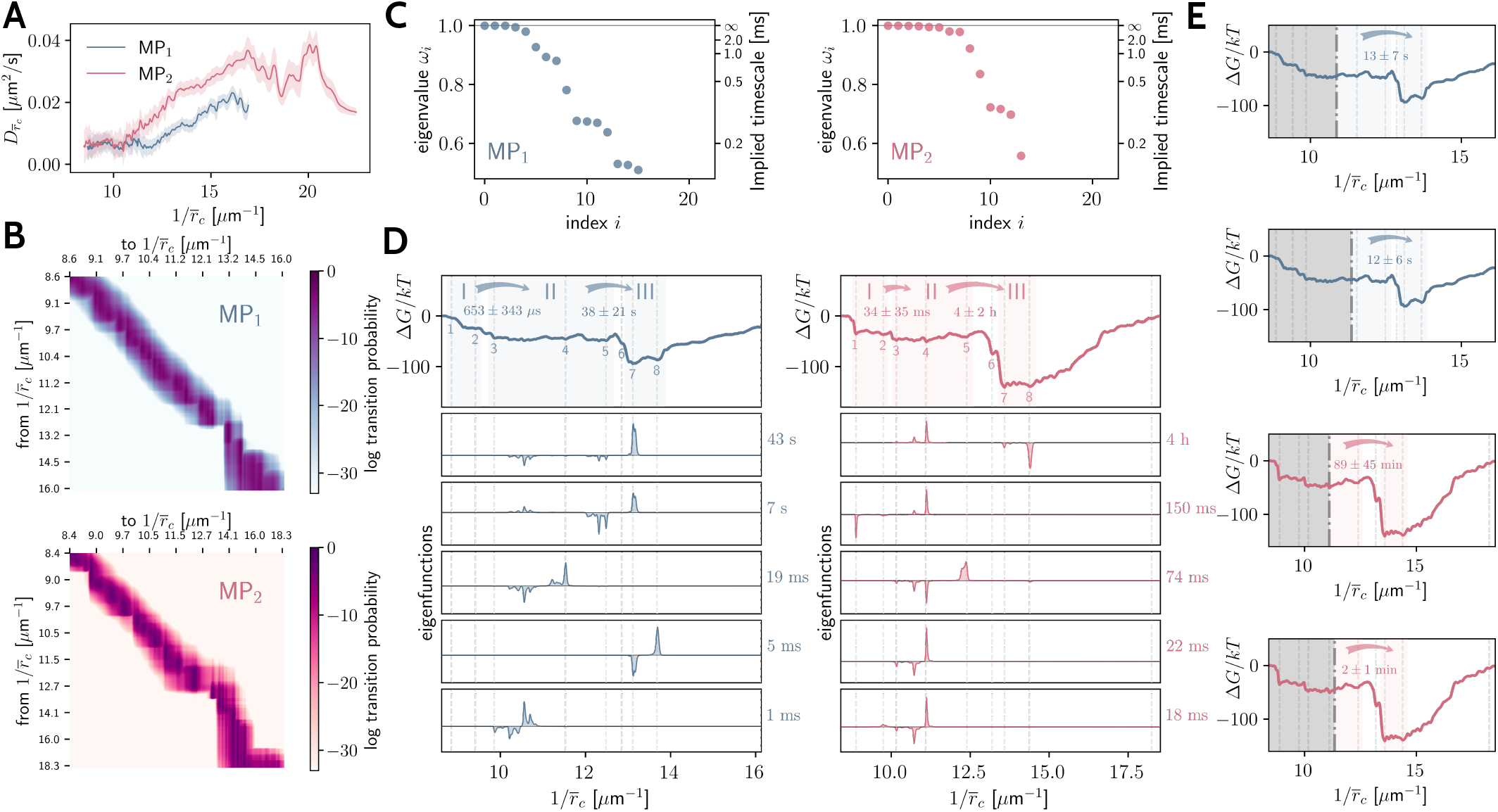
Kinetics of spontaneous membrane invagination by peripheral proteins: **(A)** Diffusion coefficient associated with the random walk in the reaction coordinate space for the two protein types. **(B)** Logarithmic colormap-representation of the transition matrix **T**, which is the discretized transfer operator for Brownian dynamics traversing the free energy landscape. Each component *T*_*ij*_ is labeled based on 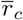 values between which the transition probability is presented. **(C)** Spectrum of eigenvalues of the transition matrix. The implied timescales corresponding to each eigenvalu can be discerned from the secondary axis. **(D)** A selection of left eigenfunctions of the discretized transfer operator, accompanied by their pertinent timescales. For reference, the upper panels replicate the free energy profiles with the enumerated states (marked as well by dashed vertical lines) coincident with those depicted in Fig. 2. Also included are the macrostates I, II, and III (shaded regions in upper panels) and the mean and standard deviation of first passage time between them. **(E)** The II ***→*** III transition kinetics when a semi-reflective boundary is introduced in the middle of macrostate II (at the position of the dot-dashed line), limiting access to states within the gray-shaded region.

Both MP_1_ and MP_2_ showcase similar diffusion coefficients, 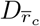, in dispersed states (Fig. 3A, smaller values of 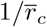). This behavior aligns with expectation, since in these states, both protein species, that have similar individual diffusion coefficients, diffuse relatively freely, devoid of effects introduced by crowding, interactions, and membrane deformations. As protein particles commence aggregation, disparities emerge. General dependence of the diffusion coefficient on 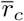 can be attributed to geometric considerations; individual protein motions substantially influence 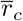 when particles cluster near the center. In such a localized cluster, diffusion of each protein can shift 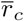 more drastically than when the particles are scattered over a large area. For MP_1_, which forms a dense, shallow pit, the significant crowding subdues this effect. In contrast, MP_2_ particles, which induce a sparser deep invagination (as seen in Fig. 2D), exhibit elevated mobility (Fig. 3A).

The next step is to build the transition matrix **T**, which is the discretized representation of the transfer operator, in the sense that each component *T*_*ij*_ is the transition probability from state *i* to state *j* during one transition step (Methods section “Kinetics”). We adopted a lag-time of 0.1 ms for estimating transition matrices. This selection stemmed from the time-correlation analysis depicted in Fig. 1D.

Comparing components of transition matrices when visualized on the same scale for both proteins (Fig. 3B), reveals that with MP_1_, the transition matrix is predominantly smooth around its diagonal, with the most formidable free energy barriers manifesting as pronounced dents. On the other hand, the MP_2_ proteins’ transition matrix has a more distinct block pattern, suggesting persistent metastable states along the reaction coordinate.

The spectrum of eigenvalues of the transfer operator for both proteins unveils the existence of several slow processes (Fig. 3C, note the cluster of eigenvalues nearing one). The transition pattern corresponding to a specific eigenvalue can be discerned by examining its associated eigenfunction. These eigenfunctions offer insight into the probability flow along the reaction coordinate [56]. Here we have inspected the left eigenvectors of the transition matrix, which correspond to transition probabilities weighted with the stationary distribution (Fig. 3D).

For both proteins, the eigenfunctions corresponding to slowest processes align with the dynamics where a pit is successfully instantiated. For MP_1_, transitions from states in either of 3 *−* 4 or 4 *−* 5 intervals to the basin at 7 occur rather fast, respectively requiring 43 s or 7 s (Fig. 3D). The rest of transitions for MP_1_ are in milliseconds range. On the other hand, the prediction for MP_2_ proteins is an initial, slow-paced formation of a vast protein cluster on the membrane, which occurs on a 100 ms timescale (Fig. 3D). The sluggishness of this process arises from the minuscule 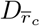 in the dispersed state as well as the metastability of small clusters at states around 1 and 4. As we previously documented, protein aggregation naturally advances toward creation of not one, but several clusters that experience pairwise repulsion [35]. This observation potentially elucidates why a singular central cluster’s inception is a slow event with MP_2_ proteins. The most drastic observation here is that for MP_2_ proteins, the estimated transition time to the initiating the invagination is roughly 4 h (Fig. 3D). We will further investigate this overshadowing timescale.

Using the transition matrix of the described finite Markov chain, we can also estimate the mean and variance of the first hitting (passage) times for coarsegrained sets of states (Methods section “Kinetics”). For the two proteins, we have selected three such clustered macrostates, shown with roman numerals in Fig. 3D – loosely corresponding to scattered (I), aggregated (II), and invaginated (III) configurations.

For MP_1_ proteins, both I ***→*** II and II ***→*** III transitions are much faster, with scattered to aggregated happening on sub-millisecond, and aggregated to invaginated on subminute times (Fig. 3D). On the other hand, MP_2_ proteins undergo a much slower transition, where tens of milliseconds is needed for aggregation and it can take up to hours for the invagination to spontaneously occur (Fig. 3D).

The timescales we have determined warrant further examination. A significant advantage of our approach is its holistic nature: rather than confining transitions to single-barrier crossings similar to Kramer’s model [57], our method embraces global transitions. Consider the II ***→*** III transition with MP_2_ proteins (Fig. 3D): it’s not merely the barrier height between states 5 and 6 that dictates the mean first passage time. Instead, it’s the presence of a profound basin within state II, centered around state 4. This nuance is echoed in the eigenfunctions linked to the slowest processes, which emphasize the influence of the distribution proximate to state 4 (Fig. 3D).

To validate this, we restricted set II to a smaller cluster near state 5 and still identified comparable extended timescales. To rectify the influence of the basin at state 4, we conducted our kinetics analysis, this time implementing a semi-reflective barrier along 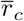 within macrostate II. This action effectively segments the 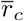 space, restricting transitions (refer to Fig. 3E). While the repercussions for MP_1_ are marginal, the kinetics of invagination for MP_2_ undergo a significant transformation,making the process potentially achievable within mere minutes.

## DISCUSSION

We have developed a particle-based membrane model wherein representative particles act as flexible, curvature-inducing proteins. The model successfully reproduces membrane invaginations in steered simulations (Fig. 1). While similar invaginations have been previously observed in dynamically triangulated membrane models using Monte Carlo simulations [26, 39], our approach offers distinguishing features: (i) our membrane and protein models contain length-scales in the form of membrane thickness and protein geometry [35], which affect the invagination morphology, and furthermore impose dynamical effects via the hydrodynamic coupling method [34]. (ii) our model inherently captures the long-range effects of curvatures induced by peripheral proteins, resulting in the emergence of membrane-mediated effects, and negating the need for their explicit enforcement [35, 37]. (iii) our use of steered molecular dynamics provides insights into the dynamics of protein clustering on the membrane and the evolution of membrane invaginations. We therefore believe our results to be better candidates for complementing experimental observations.

The free energy profiles derived for the two proteins consistently showed an initial drop, indicative of formation of small but stable clusters. This was succeeded by energy barriers impeding large-scale aggregation and pit emergence. However, MP_2_ proteins, inducing pronounced membrane curvatures, exhibited a rougher landscape dotted with prominent metastable states. These proteins also successfully transformed pits into membrane invaginations (Fig. 2 and Supplementary Movies). From the theoretical standpoint on membrane-mediated interactions, pairs of proteins that induce same-sign isotropic curvatures should repel [58, 59]. On the other hand, stiff-binding proteins that locally suppress membrane fluctuations experience attractive interactions of entropic nature [37] – which has been dubbed the thermal Casimir effect [59–61] Finally, when proteins are in close vicinity, and have introduced large contact angles, stronger attraction can be achieved via a mechanism not dissimilar to the “cheerios effect” [62, 63]. All these pictures are somewhat idealized models pertaining to dilute systems of proteins and neglecting multi-body effects [64, 65]. In a dynamic and dense assembly of stiff-binding membrane-bending proteins, significant forces of varying nature may be present [37, 66]. Metastable structures, forming dynamically through the stochastic motion of proteins and membrane, can thus determine the global trend.

Previously, we had exhaustively validated the kinetics of the current model at its scale of application [33– 35]. We also took great care in establishing the protocol for the steered simulations in this work. This lends credence to the kinetic predictions of the model, and allows us to compare the resulting timescales with the experimental observations. We may compare the timescale obtained here with the time it took STxB proteins to induce tubules when incubated with giant unilamellar vesicles (GUV’s) [13]. In these experiments, tubulation had been observed after several minutes. On the surface, it might seem that we have made a paradoxical observation: With MP_1_ proteins, the kinetics of pit formation is on a comparable timescale. However, they ultimately fail to induce a deep invagination. On the other hand, the more successful MP_2_ proteins apparently take several hours to spontaneously invaginate the membrane (Fig. 3).

In general, this dilemma is worth a more detailed study, especially considering uncertainties in the estimated free energies and diffusion coefficients. Nevertheless, we observed that limiting the system dynamics slightly away from one dominant metastable state can significantly shift the kinetics toward the expected range (Fig. 3E). In practice, such an effect may arise from (i) higher surface concentration of proteins, or alternatively, global fluctuations in surface concentration resulting in locally dense pockets, (ii) having an open system with the possibility of protein adsorption. We have modeled the invagination process as occurring under the influence of proteins already bound to the membrane, with an initial scattered configuration. Furthermore, we have simulated a membrane patch of limited size, with a specific path for steered simulations, both of which constrain the surface density fluctuations. Thus, the limitation that we artificially introduced over 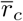 can actually point to different scenarios where *r*_*c*_ around one invagination site has a natural upper bound due to surface concentrations.

The recruitment and binding of toxins such as Shiga and Cholera with pentameric membrane-binding sub-units is in itself a process worthy of detailed study [67]. It is imaginable that including a reservoir of proteins ready to be recruited [68] or performing grand canonical simulations can lead to smoother energy landscapes and faster kinetics.

## CONCLUSION

This work elucidates the intricacies of membrane invagination driven by curvature-inducing flexible proteins. By leveraging the Jarzynski equation, we delineated the free energy landscape of the process, comparing two distinct protein types, MP_1_ and MP_2_. We revealed how disparities in their local curvature distributions led to variances in global conformations, energy landscapes, and kinetics.

Our methodology could potentially be adapted to analogous systems, notably those involving BAR-domain proteins with anisotropic curvatures. Furthermore, our free energy estimation procedure may be utilized to shed light on mechanisms underpinning processes like clathrin-independent endocytosis [69] or endoplasmic reticulum reorganization [70].

There are multiple avenues for refinement, including employing more realistic peripheral protein models, accounting for membrane tension, and considering the influence of cytoskeletal structures. Our innovative tools and methodologies present exciting possibilities for the future, enabling a deeper comprehension of the membrane-protein dynamic.

Ultimately, we envision this work’s most ambitious application as a foundation for engineering endocytosis, propelling the use of proteins like STxB for targeted drug and antigen delivery.

## METHODS

### Free energy estimation

We have used the Jarzynski equality for free energy estimation in steered simulations. This equation relates the free energy difference between two states 1 and 2 Δ*G* = *G*_2_ *− G*_1_, to the non-equilibrium work, ***δ****W*_1 ***→***2_, performed in bringing the system from 1 to 2 [48],

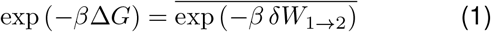

where *β* = 1*/kT*, and the average is over an ensemble of transition paths. The Jarzynski equality is valid as long as the probability distribution of microstates at the starting state is given by the Boltzmann distribution exp [− *β*ℌ_0_ (**z**)] */Z*, where we use the phase vector **z** to simultaneously describe the configuration and momenta of the system, and *Z* denotes the partition function with *G* = *− kT* ln *Z*.

The external work resulting from finite-time evolution of the system along the trajectory *λ* (*t*), where *λ* is a (macroscopic) reaction coordinate, is considered as [48, 49]:

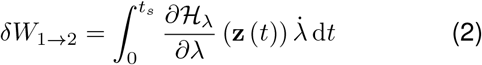

where *t*_*s*_ is called the switching time.

We have done our steered simulations in an isothermal-isobaric ensemble, where barostat coupling to a setpoint pressure *p*_*XY*_ happens in the in-plane *X* and *Y* directions. The Hamiltonian that would thus describe the equilibrium distribution at any point along the trajectory *λ* (*t*) is [71],

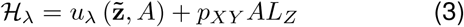

where *u*_*λ*_ is the energy of the system, 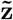 is the phase vector scaled down by the in-plane characteristic dimension 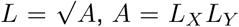 is the projected area of the membrane patch, and *L*_*Z*_ is the box dimension in the direction perpendicular to the plane of the membrane. As the barostat scales both in-plane dimensions in a coupled manner, the projected area is enough to related 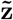 to z. Thus, we have,

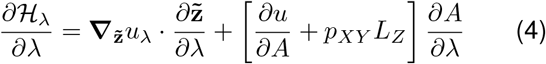

In our case the reaction coordinate *λ* is simply given as,

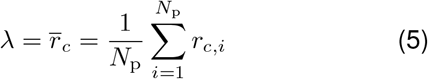

where *r*_*c,i*_ is the two-dimensional radial distance of the *i*-th protein particle from the central axis of the simulation box, and *N*_p_ is the total number of proteins on the membrane surface. It is to be noted that wherever we presented the results as a function of 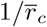, this was done after the fact, and all calculations were done originally with 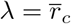.

Limiting the following derivations to the configurational part of 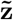 denoted as 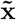, and setting the origin in the middle of the simulation box, coinciding with the initial plane of the membrane, we note that 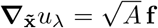 with f being the vector of external centripetal forces.

The gradient 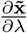 is non-zero only for the subset of particles contributing to the reaction coordinate, *λ*. However, as is usual with collective variables, this gradient is not uniquely defined, and only needs to satisfy the orthogonality condition 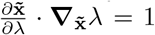 [72]. Based on eq. (5), we find,

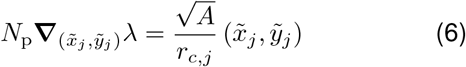

for the *j*-th protein particle, where 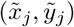 means either of its in-plane coordinates,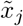 or 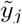. We can argue based on symmetry that there should be no distinction between particle positions contributing to the eq. (5). Thus, we consider,

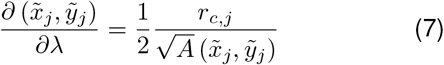

leading to the first term in the right-hand side of eq. (4) becoming − *N*_p_ |**f**|.

We can further substitute 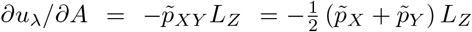 and *p*_*XY*_ = 0 (notice the difference between instantaneous values of pressure, 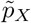 and 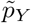 versus the setpoint in-plane pressure *p*_*XY*_ assigned to the mechanical bath via the barostat).

In order to obtain *∂A/∂λ*, it is beneficial to consider the simultaneous scaling of the simulation box in the *X* and *Y* direction with an instantaneous “strain”, *ϵ*, such that *L*_*X*_ = (1 + *ϵ*) *L*_*X*,0_ and *L*_*Y*_ = (1 + *ϵ*) *L*_*Y*,0_, in which *L*_*X*,0_ and *L*_*Y*,0_ are the box dimensions before the scaling. We would thus have, *∂λ/∂ϵ* = *λ*_0_ and *∂A/∂ϵ* = 2 (1 + *ϵ*) *A*_0_, where similarly, *λ*_0_ and *A*_0_ correspond to the state before the scaling. This results in 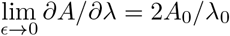.

To employ eq. (1) and obtain mean free energy differences and their uncertainties, we used the following procedure:

1. To ascertain that the trajectories originate from an equilibrium distribution, we accumulated all the trajectories pertaining to initial states into one ensemble, for which a probability distribution over the reaction coordinate was established via histogramming.
2. Each whole trajectory is then reweighted based on its occurrence probability at the onset of external loading, via Markov chain Monte Carlo sampling of trajectory replicas.
3. We set out to estimate the external work from eq.(2) along each simulation trajectory. As each trajectory may visit macrostates along *λ* more than once, all the estimates of the gradient of the Hamiltonian are gathered in global bins of *λ*.
4. An ensemble of external work samples are produced via bootstrapping through which samples of the mean of the gathered Hamiltonian gradients are integrated. The procedure is repeated for each trajectory, and leads to the results shown in Fig. 2A).
5. The Jarzynski equality is finally applied to the obtained ensemble of external work over global *λ* bins. Bootstrapping is again used over samples the external work up to each *λ* bin, yielding the uncertainty in the average of the exponential in eq. (1).

### Kinetics

Given the free energy profile as a function of the reaction coordinate *λ*, one simple approach to estimating the kinetics of spontaneous processes is to consider a diffusion process on this one-dimensional energy landscape [73]. To estimate the timescales of slow processes in such a description, a reliable method is to use the transfer operator. Taking the time-dependent probability density *P*_*t*_ (*λ*) over states along the reaction coordinate *λ*, and its weighted counterpart 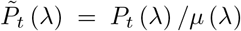 where *μ* (*λ*) is the equilibrium (stationary) probability density, the transfer operator _*τ*_ with the lag-time *τ* is defined as [56, 74],

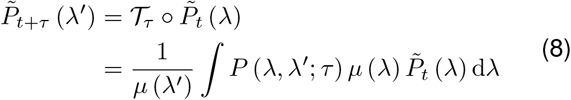

where *P* (*λ, λ1*; *τ*) is the transition probability density between states *λ* and *λ1*.

Eigenvalues of the transfer operator (*ω*_*i*_’s) are related to the timescales of transitions as *t*_*i*_ = *− τ/* ln *ω*_*i*_ [56, 75]. Assuming Brownian dynamics in the reaction-coordinate space, we can estimate discretized transition probabilities directly [76],

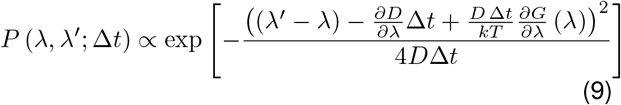

where *∂G − /∂λ* is the mean force.

To build the transition matrix **T**, a spatial grid of 800 bins is considered over the range of 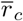. The timestep of the Brownian motion, Δ*t*, is estimated based on the minimum value of 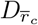, and the size of one grid cell, such that the timestep accommodates transitions to neighboring cells. We derived values of Δ*t* = 565 ns for MP_1_ and Δ*t* = 982 ns for MP_2_. The lag-time is hence divided into ⌈ *τ/* (2Δ*t*) ⌈ sub-steps, and the transition matrix for the whole lag-time is simply obtained as the product of transition matrices of the sub-steps.

In order to estimate mean and variance of first passage times from the transition matrix **T**, we start by changing destination states to absorbing. This results in the following partitioning of the transition matrix,

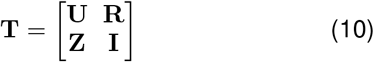

where **U** contains the transition probabilities within non-absorbing states, **R** components are the transition probabilities to the absorbing states, **Z** is a matrix of zeros, and **I** is the identity matrix. With these notations, the mean and variance of first passage times, *n* for a process starting with an initial probability vector of **p** are found via,

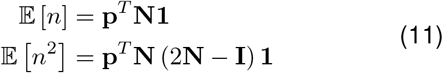

where *n* is the random variable describing discrete number of steps until the process is absorbed, and **N** = (**U***−* **I**)^*−*1^ is the fundamental matrix of the Markov chain, and **1** is a column vector of ones. For the initial probability vector, we use a local distribution over the starting set that is proportional to the global stationary distribution *μ* (*λ*).

### Simulations

The membrane model used for the simulations presented here consists of a bilayer of particles, each linked to its nearest neighbors through Morse-type bonds. In addition, each pair of opposing particle from the two leaflets is harmonically bonded. Harmonic angle-bending potentials are used to constrain the rotational motion of in-plane bonds relative to out-of-plane ones. The force field parameters are derived by optimizing the energy density of the discrete model against the continuum elastic model of the membrane. Bond-flipping Monte Carlo moves are incorporated to simulate inplane fluidity and permit lateral particle diffusion [29].

More recently, we showed how peripheral proteins can be incorporated in the model via force field masking [35, 37]. The parameter-space optimization method is adapted by including a continuum representation of the protein on top of the membrane. The protein possesses an intrinsic curvature and finite flexibility, and thus contributes to the total elastic energy. For the two proteins studied here, we have used a geometric representation similar to the membrane-binding subunit of Shiga toxin, STxB. An isotropic elastic modulus is assigned to the protein, which is 100 MPa for MP_1_ and 200 MPa for MP_2_ (for full details see [35] and its supplementary information). In the end, what can be considered equivalent to the protein model in our simulations is the distribution of local curvatures shown in Fig. 1B.

Flat membrane patches of 325 nm lateral size with a number of bound peripheral proteins randomly scattered on their surface are used as the initial configuration of each simulation. For each of the two protein types, several realizations of the steered simulations that start from different initial configurations are performed. External forces applied on protein particles are updated in each time step to be continually directed toward the central axis of the simulation box (Fig. 1A).

During each simulation, particle positions have been propagated using anisotropic over-damped Langevin dynamics that includes hydrodynamic interactions between nearest neighbors [33, 34]. Water, with the dynamic viscosity of 0.7 mPa s, has been used as the surrounding fluid environment. Simulation box is laterally coupled to a zero-pressure bath via a stochastic Langevin-piston barostat [77, 78]. All simulations have been performed at constant 310 K temperature.

Simulations are performed using an in-house special-purpose software. Simulation snapshots and movies are visualized using the Visual Molecular Dynamics (VMD) software [79]. All the analysis presented are performed with in-house Python codes, using Numpy [80] and Scipy [81] numerical packages.

## AUTHOR CONTRIBUTIONS

The author developed the model and the theory, performed the simulations and analyzes, and prepared the manuscript.

## FUNDING

This research has been funded by Deutsche Forschungsgemeinschaft (DFG) through grants SFB 958/A04 and SFB 1114/C03.

### ACKNOWLEDGMENTS

The author is especially grateful to Frank Noé (FU Berlin) for constant support and valuable comments and discussions.

## DATA AVAILABILITY STATEMENT

The data that support the findings of this study are available from the author upon reasonable request.

